# The sensor hub for detecting the developmental characteristics in reading in children on a white vs. coloured background/coloured overlays

**DOI:** 10.1101/2020.08.04.235846

**Authors:** Tamara Jakovljević, Milica Janković, Andrej Savić, Ivan Soldatović, Petar Todorović, Tadeja Jere Jakulin, Gregor Papa, Vanja Ković

## Abstract

The study investigated the influence of white vs 12 background and overlay colours on the reading process in school age children. Previous research reported that colours could affect reading skills as an important factor of the emotional and physiological state of the body and that reading is one of the most important processes in the maturation of children. The aim of the study was to assess developmental differences between second and third grade students of elementary school and to evaluate differences in electroencephalography (EEG), ocular, electrodermal activities (EDA) and heart rate variability (HRV). In the experiment, the responses of 24 children (12 second and 12 third grade students) to different background and overlay colours were summarized using EEG, eye tracking, EDA and HRV signals. Our findings showed a decreasing trend with age regarding EEG power bands (Alpha, Beta, Delta, Theta) and lower scores of reading duration and eye-tracking measures in younger children compared to older children. As shown in the results, HRV parameters showed higher scores in 12 background and overlay colours among second than third grade students which is linearly correlated to the level of stress and readable from EDA measures as well. The existing study showed the calming effect on second graders in turquoise and blue background colours. Considering other colours separately for each parameter, we assumed that there are no systematic differences in Reading duration, EEG power band, Eye-tracking and EDA measures.

## Introduction

Learning to read is a complex process involving both perception and cognition, via integration of visual and auditory information processing and memory, attention and language skills [1]. Therefore, reading is a taught skill depending upon a range of perceptual processes and cognitive abilities affecting learning over time and across development [2–4] . One of the key objectives of early education is the learning of reading, and a number of research studies have explored the process of reading skills acquisition in children [2,5–7]. Depending on their individual set of underlying abilities, children will have different developmental profiles of reading skill obtainment, while weaknesses in some abilities may cause reading impairments over time. Individual differences in learning of reading may originate from biological and environmental factors, shaping the development of brain systems involved in the reading process [8]. The current knowledge emphasizes the importance of identifying and treating reading difficulties as early as possible since they may impair academic achievements and increase the risk of social, emotional and mental health problems in children. More specifically, poor reading skills are shown to be associated with increased risks for school dropout, attempted suicide, incarceration, anxiety, depression, and low self-concept [9] ,

There is some limited evidence that colours may impact the reading process, specifically with early school-age children, and those with reading disabilities [10,11]. Going back a few decades [12,13], it has been shown that the role of colours in reading dates back to 1958. Jansky [14] reported the case of a student with a reading deficit who was able to recognize words printed on yellow paper, but unable to recognize words printed on white paper. Previous studies considered the influence of background-, text- or overlay-colour on the actual reading process in children [13,15,16]. While more recent studies have shown that colours do not influence the reading process and that this could be a placebo [17], others have found that colours may be particularly effective for early readers in school-age children [18].

As reading involves sensory integration, attention and memory, those processes may be reflected in the psycho-physiological states of the individual engaged in the reading task. Those states are a result of underlying neural and physiological processes, which are measurable and quantifiable by different biosignal modalities. The goal of this study was to employ multimodal sensor measurements to examine the influence of colour of the content on the reading task in children at different developmental stages. More specifically, we have employed measurements of electroencephalography (EEG), eye-tracking, electrodermal activity (EDA) and heart rate variability (HRV) to assess the influence of background and overlay colour on reading performance in second and third grade students of elementary school. We aimed to address the mechanisms of colour influence on the reading process through electrophysiological correlates of the reader's state, while taking into account the developmental aspect of reading acquisition.

This study sheds light on underlying neural, physiological and behavioural processes accompanying the reading task in children. Moreover, this is an initial step towards the possibility of including colour into reading content to improve reading skills in children at different developmental stages.

## Method

### Participants

Twenty-five healthy participants were randomly chosen from a single class of the second and third grades of the elementary school “Drinka Pavlović” in Belgrade respectively. The participants ages ranged from 8 to 9 years. Inclusion criteria were that children have no reading and learning disabilities, attention disorders, have normal or corrected to normal vision. Only one child was excluded from the analysis due to the large artefacts in the acquired signals and his data were not used in the statistical analysis.

All the subjects underwent the same experimental conditions and participated individually in the small school classroom during the regular time period of the daily school classes. Each child received a short instruction about the experiment setup and design. After the experiment, every child received a small present and diploma for participation in the experiment process.

The ethical committee of the Psychology Department of the University of Niš approved the experimental procedure.

### Experiment setup

During the experiment, each participant was sitting on a chair at a table in the front of a computer monitor and keyboard. At the beginning of the experiment, participants got the instruction to read quietly for themselves the text from the stimuli presentation shown on the computer monitor and to press the space button for the next slide of stimuli presentation. The experiment was repeated in one pseudo randomization of colour background/overlay order starting always with a referent slide (black text on white background). Details about the “experiment” content are explained in the Section Experiment Design.

During the reading process, physiological data were acquired using a sensor hub composed of an eye-tracking system and a portable multimodal EEG/ECG/EDA system, Fig. 1. Two laptops were used for data real-time monitoring and storage: one laptop for eye movement monitoring (with additional external computer monitor and keyboard in front of the participant) and another for EEG/ECG/EDA monitoring.

**Figure 1.**
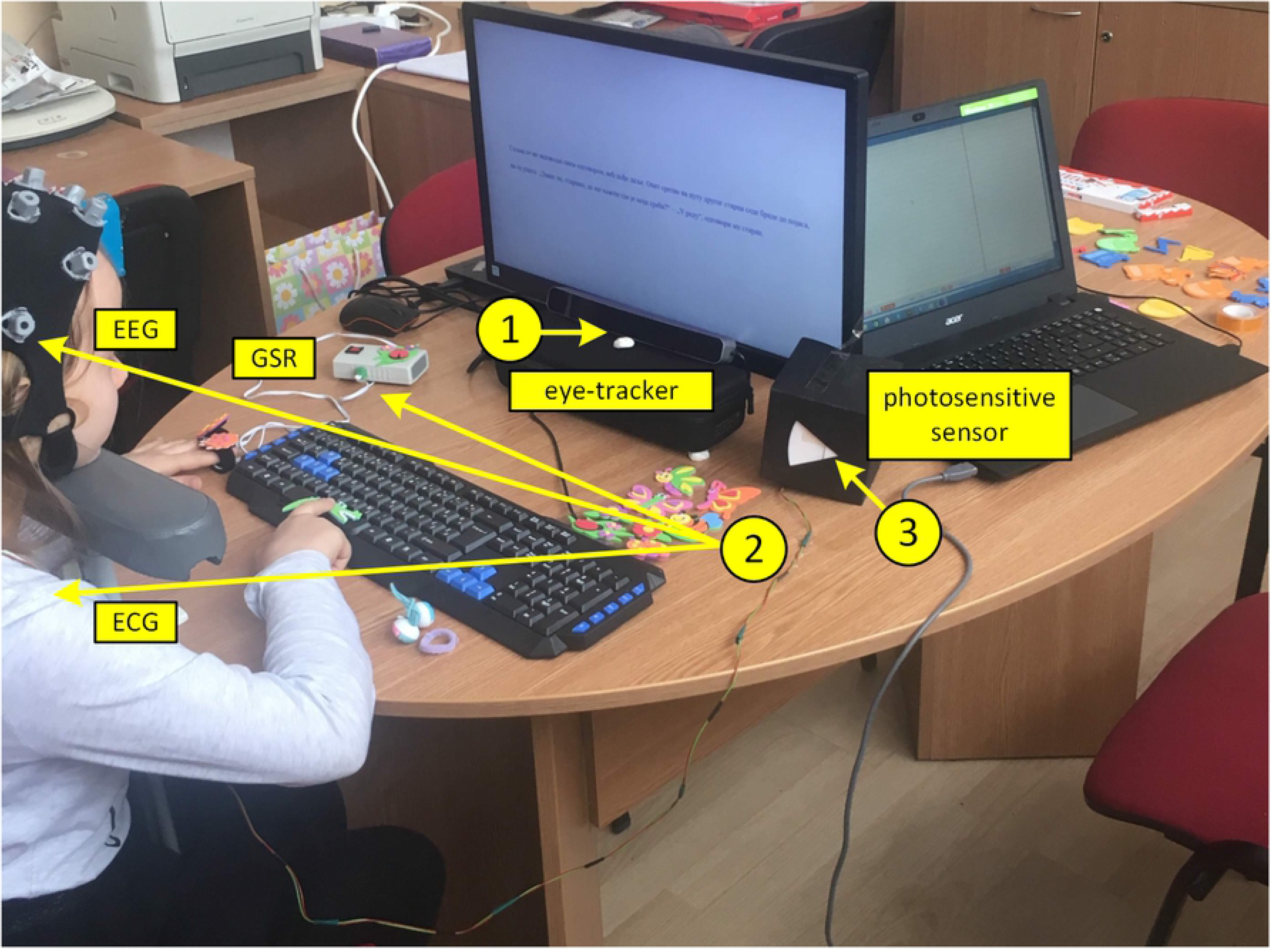
Experiment setup: 1) Eye-tracking system 2) Portable multimodal EEG/ECG/EDA system, 3) Photosensitive sensor for synchronization of multimodal EEG/ECG/EDA system and eye-tracking system.

An SMI RED-m 120-Hz portable remote eye tracker (https://www.smivision.com) was placed below the computer monitor in front of the participant, and it was fixed in place to keep it from accidentally moving. An adjustable chin-rest was used to ensure the same distance from the monitor and table for each participant (a chinrest was 57 cm away from the eye-tracking sensor and 16 cm above the table) [19]. The SMI software was used for stimuli presentation (Experiment Centre 3.7) and data collection (iView RED-m).

The EEG and ECG signals were recorded using a mobile 24-channel EEG amplifier (SMARTING, mBrainTrain, Belgrade, Serbia) wirelessly communicating with a laptop via Bluetooth. Twenty-two monopolar EEG channels of Greentek Gelfree-S3 cap (10/20 locations: Fp1, Fp2, F3, F4, C3, C4, P3, P4, O1, O2, F7, F8, T7, T8, P7, P8, Fz, Cz, Pz, AFz, CPz, POz) were recorded. The ground was located at FPz and FCz was used as the reference site. One channel of the amplifier was connected to the surface SKINTACT ECG electrode placed in the left chest region, over the heart, to record ECG signal as a reference for heartbeat detection. EEG and ECG signals were acquired with 24-bit resolution and 250 Hz sampling rate. The skin-electrode impedances were below the manufacturer recommended value of 1 kOhm, prior to the tests.

One channel of the amplifier was used for the synchronization of electrophysiological recordings and eye tracking data. A small photosensitive sensor registering the changes on the screen after each slide was sused with the changes of the black and white screen (200 ms each) and sending a trigger for synchronization of multimodal EEG/ECG/EDA system and eye tracking system, for each event (slide).

We used a research prototype for galvanic skin response (electrodermal activity, EDA) recording [20] that communicates with a laptop via Bluetooth. The sampling rate for EDA data was 40 Hz. SMARTING application has Lab Streaming Layer (LSL) compatibility which enabled synchronization between EDA and EEG/ECG data within a single file of XDF format.

### Experiment design

#### Stimuli

Participants read a story on the computer monitor. The story was at an adequate level for the second/third-grade of elementary school and selected from the school literature of the Serbian language course. Participants were unfamiliar with the text used in the study.

The story “St Sava and the villager without luck” was split into 13 paragraphs: the 1st slide as a referent one - white background with black letters, then 6 slides with black letters on red, blue, yellow, orange, purple and turquoise background and the next 6 slides in the overlay manner which looks like covering black text on a white background with a coloured foil (calculated by the algorithm described in the Section Colour calculation).

The experiment started with calibration and validation method would stop on the black slide so the researcher had time to launch multimodal EEG/ECG/EDA system and eye-tracking system for data acquisition. Next, on the researcher's instruction, the child would press the space button on the keyboard and the first slide with the text appeared on the computer monitor. The participant read text to themselves and then pressed the space button for the next slide to continue the text.

After finishing the test, the researcher checked with each child the level of understanding of the story with questions recommended in the literature after the story for exercise.

#### Colour calculation

All colours (colour shades) used for designing the slides (stimuli) were defined within the RGB colour model and each individual colour was expressed as a RGB triplet [R,G,B], where the value of each additive primary colour component can vary from 0 to 255. List of background shades in slides with coloured background (and black text) with associated numerical values of their RGB triplet are: red (“red”, [225,0,0]), blue (“blue”, [0,0,225]), yellow (“yellow”, [225,225,0]), orange (“orange”, [225,128,0], purple (“purple”, [225,0,128]) and turquoise (“turquoise”, [0,225,225]. White and black shades were defined by triplets [225,225,225] and [0,0,0], respectively. RGB components of the background and text in the slides with "overlay effect" were calculated according to the following formula:

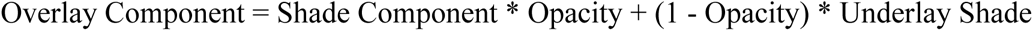

where, Opacity value was set to 0.5, Shade Component was selected from one of the previously listed background colour shades, and Underlay Shade value in case of text being 0 for black and in case of white background being 255.

The resulting RGB triplets for shades of the text and background for slides with overlay effect were: overlay red (“red O”, text - [128,0,0], background -[225,128,128]), overlay blue (“blue O”, text - [0,0,128], background -[128,128,225]), overlay yellow (“yellow O”, text - [128,128,0], background -[225,225,128]), overlay orange (“orange O”, text - [128,64,0], background - [225,192,128]), overlay purple (“purple O”, text - [128,0,64], background -[225,128,192]) and overlay turquoise (“turquoise O”, text - [0,128,128], background -[128,225,225]).

#### Data processing

Eye tracking data analysis and visualization was performed using BeGaze 3.7 software. The selected eye-tracking parameters were: fixation count, fixation frequency (count/second), fixation duration total (ms), fixation duration average (ms), saccade count, saccade frequency (count/second), saccade duration total (ms), saccade duration average (ms).

EEG/ECG/EDA data were analysed using Matlab ver. 8.5 (Mathworks, USA) in the manner described below.

For each subject and electrode site, EEG signal was processed in order to calculate the band power in 5 predefined frequency bands: delta (0.5 - 4 Hz), theta (4 - 7 Hz), alpha (7 - 13 Hz), beta (15 - 40 Hz) and whole range (0.5 - 40 Hz). Median value of EEG band power (for each frequency band) was determined for 13-time epochs coinciding with reading of the content of each presented slide. Median EEG power in each frequency band was calculated by raw continuous signal band-pass filtering (4th order Butterworth filter with cut-off frequencies defined by individual band's frequency range), squaring, segmenting into 13 epochs (determined as time intervals between each slide's onset and offset) and median averaging to a single power value over signal samples of each epoch (i.e. over each slide's duration). Applied processing resulted in the 65 median power values (i.e. 13 slides x 5 frequency bands) for each EEG channel of each subject. Median power calculation was applied since it is less likely to be affected by outliers in the EEG power samples occurring with temporary movement artefacts than the mean power.

Heart activity signal was band-pass filtered using a FIR filter in the range 1-45 Hz (750 points), after which the Pan-Tompkins algorithm [21] for extraction of heart activity beats was applied. Beat-to-beat intervals (BBI, the time between two successive heart activity peaks) were calculated. Heart rate variability (HRV) parameters [22] were calculated from BBIs for 13-time epochs coinciding with reading of the content of each presented slide, Table 1. Applied processing resulted in the 14 HRV parameter values in one subject for each of the 13 slides.

**Table 1.** HRV parameters

| Parameter | Unit | Description |
| --- | --- | --- |
| **Time domain parameters** |  |  |
| Mean RR | ms | Mean value of BBIs |
| SDNN | ms | Standard deviation of normal BBIs |
| Mean HR | beats/min | Mean value of heart rate |
| STD HR | beats/min | Standard deviation of heart rate |
| CVRR=SDNN/Mean RR | n.u. | Coefficient of variance of normal BBIs |
| RMSSD | ms | Root mean square of differences of successive BBIs |
| NN50 | beats | Number of successive BBIs that varied more than 50 ms |
| pNN50 | % | Percentage of successive BBIs that differ more than 50 ms |

Mean value of EDA data was calculated as a representative value of electrodermal activity for 13-time epochs coinciding with reading of the content of each presented slide. Applied processing resulted in the 13 mean values for each subject (one mean EDA value per slide).

### Statistical methodology

Results are presented as count (%), means ± standard deviation or depending on data type and distribution. Groups are compared using parametric test, independent samples t test. All p values less than 0.05 were considered significant. All data were analysed using SPSS 20.0 (IBM Corp. Released 2011. IBM SPSS Statistics for Windows, Version 20.0. Armonk, NY: IBM Corp.) and R 3.4.2. (R Core Team (2017). R: A language and environment for statistical computing. R Foundation for Statistical Computing, Vienna, Austria. URL https://www.R-project.org/.).

## Results

### White (default) background - reading results

Grade comparisons (second vs. third) regarding the examined parameters for white colour only are presented in Table 2. A significant difference has been obtained regarding EEG frequency bands (Alpha, Beta and Theta) and ECG parameters (SDNN, CVRR and STD HR). In all other parameters we observed no significant difference between second and third graders. All EEG power bands are higher in the second grade group (except Delta band), compared to third graders. An opposite trend is shown in SDNN, CVRR and STD HR parameters whereby third graders achieved higher scores in comparison to second graders.

**Table 2.** Reading duration, EEG, Eye tracking, EDA and HRV parameters in second and third grade - significant p values are marked as bold

| Parameters | Grade |  | p value* |
| --- | --- | --- | --- |
|  | Second (n=12) | Third (n=12) |  |
| Reading duration |  |  |  |
| RD (s) | 43.38 ± 25.32 | 49.07 ± 27.63 | 0.628 |
| EEG parameters (median power band) |  |  |  |
| Alpha | 13.19 ± 5.15 | 5.63 ± 4.28 | **0.001** |
| Beta | 6.05 ± 2.81 | 3.33 ± 2.40 | **0.018** |
| Delta | 64.45 ± 20.47 | 71.85 ± 63.54 | 0.707 |
| Theta | 16.07 ± 7.14 | 8.35 ± 7.67 | **0.018** |
| Whole Range | 113.3 ± 38.5 | 97.2 ± 80.5 | 0.540 |
| Eye tracking parameters |  |  |  |
| Fixation Count | 38.25 ± 19.62 | 35.00 ± 9.66 | 0.620 |
| Fixation Frequency [count/s] | 1.01 ± 0.36 | 0.99 ± 0.67 | 0.925 |
| Fixation Duration Total [s] | 40.87 ± 25.01 | 46.22±26.32 | 0.631 |
| Fixation Duration Average [ms] | 1088.2 ± 542.9 | 1331.5 ± 730.9 | 0.381 |
| Saccade Count | 34.92 ± 19.16 | 28.70 ± 5.23 | 0.301 |
| Saccade Frequency [count/s] | 0.95 ± 0.38 | 0.86 ± 0.65 | 0.686 |
| Saccade Duration Total [ms] | 741.3±449.4 | 680.0±276.7 | 0.711 |
| Saccade Duration Average [ms] | 20.93 ± 2.76 | 23.22 ± 5.61 | 0.227 |
| EDA value |  |  |  |
| EDA (uS) | 9.41 ± 3.49 | 6.20 ± 4.38 | 0.060 |
| HRV parameters |  |  |  |
| Mean RR (ms) | 637.4 ± 59.8 | 680.5 ± 114.4 | 0.264 |
| SDNN (ms) | 35.14 ± 11.95 | 66.11 ± 46.15 | **0.043** |
| CVRR (n.u.) | 0.06±0.02 | 0.10±0.05 | **0.021** |
| Mean HR (beats/min) | 94.87 ± 8.69 | 90.26 ± 13.75 | 0.337 |
| STD HR (beats/min) | 5.12 ± 1.54 | 8.40 ± 3.85 | **0.016** |
| RMSSD (ms) | 42.50 ± 17.42 | 84.94 ± 74.74 | 0.079 |
| NN50 (beats) | 12.42±9.44 | 19.67±16.43 | 0.298 |
| pNN50 (%) | 23.40±18.34 | 36.30±25.88 | 0.236 |

\*independent sample t test

### Background and overlay colours

In the Table 2 we showed the overall results which were calculated by subtracting parameters acquired for white colour from parameters acquired for each of the background and overlay colours (normalized values). Second and third graders were then compared on each of the parameters measured in the study, namely: Reading duration, EEG, Eye tracking, EDA and HRV.

The results demonstrated that students in the second grade differed significantly from the students in the third grade consistently on a few HRV parameters, in particular: SDNN (yellow, turquoise and turquoise O), CVRR (for orange, turquoise and blue O), and STD HR (blue, yellow, turquoise, blue O, yellow O, orange O and turquoise O). Besides these differences, significant differences were found in RMSSD (for turquoise and blue O). In each of these cases second grade students scored higher (or positively) in comparison to the third grade students who scored lower (or negatively), which is marked with orange colour in Table 3. Out of all parameters in the study, only in the case of Saccade Duration Total for the blue O, the third graders scored significantly higher than the second graders, which is marked with green colour in Table 3.

**Table 3.** Differences between second and third graders on reading duration, EEG, Eye tracking, EDA and HRV parameters (normalized on white colour), (p<.05 is marked with “+”, and p<.01 is marked with “++”)

| Parameters | Normalized values |  |  |  |  |  |  |  |  |  |  |  |
| --- | --- | --- | --- | --- | --- | --- | --- | --- | --- | --- | --- | --- |
|  | r<br>e<br>d | b<br>l<br>u<br>e | y<br>e<br>l<br>l<br>o<br>w | o<br>r<br>a<br>n<br>g<br>e | p<br>u<br>r<br>p<br>l<br>e | t<br>u<br>r<br>q<br>u<br>o<br>i<br>s<br>e | r<br>e<br>d<br>O | b<br>l<br>u<br>e<br>O | y<br>e<br>l<br>l<br>o<br>w<br>O | o<br>r<br>a<br>n<br>g<br>e<br>O | p<br>u<br>r<br>p<br>l<br>e<br>O | t<br>u<br>r<br>q<br>u<br>o<br>i<br>s<br>e<br>O |
| **Reading duration** |  |  |  |  |  |  |  |  |  |  |  |  |
| RD (s) |  |  |  |  |  |  |  |  |  |  |  |  |
| **EEG parameters (median power band)** |  |  |  |  |  |  |  |  |  |  |  |  |
| Alpha |  |  |  |  |  |  |  |  |  |  |  |  |
| Beta |  |  |  |  |  |  |  |  |  |  |  |  |
| Delta |  |  |  |  |  |  |  |  |  |  |  |  |
| Theta |  |  |  |  |  |  |  |  |  |  |  |  |
| Whole Range |  |  |  |  |  |  |  |  |  |  |  |  |
| **Eye tracking parameters** |  |  |  |  |  |  |  |  |  |  |  |  |
| Fixation Count |  |  |  |  |  |  |  |  |  |  |  |  |
| Fixation Frequency [count/s] |  |  |  |  |  |  |  |  |  |  |  |  |
| Fixation Duration Total [s] |  |  |  |  |  |  |  |  |  |  |  |  |
| Fixation Duration Average [ms] |  |  |  |  |  |  |  |  |  |  |  |  |
| Saccade Count |  |  |  |  |  |  |  |  |  |  |  |  |
| Saccade Frequency [count/s] |  |  |  |  |  |  |  |  |  |  |  |  |
| Saccade Duration Total [ms] |  |  |  |  |  |  |  | - |  |  |  |  |
| Saccade Duration Average [ms] |  |  |  |  |  |  |  |  |  |  |  |  |
| **EDA value** |  |  |  |  |  |  |  |  |  |  |  |  |
| EDA (uS) |  |  |  |  |  |  |  |  |  |  |  |  |
| **HRV parameters** |  |  |  |  |  |  |  |  |  |  |  |  |
| Mean RR (ms) |  |  |  |  |  |  |  |  |  |  |  |  |
| SDNN (ms) |  |  | + |  |  | ++ |  |  |  |  |  | + |
| CVRR |  |  |  | + |  | + |  | + |  |  |  |  |
| Mean HR (beats/min) |  |  |  |  |  |  |  |  |  |  |  |  |
| STD HR (beats/min) |  | + | + |  |  | ++ |  | + | + | + |  | + |
| RMSSD (ms) |  |  |  |  |  | + |  | + |  |  |  |  |
| NN50 (beats) |  |  |  |  |  |  |  |  |  |  |  |  |
| pNN50 (%) |  |  |  |  |  |  |  |  |  |  |  |  |

In the Figure 2, normalized SDNN, CVRR and STD HR values across grades and colours are presented.

**Figure 2.**
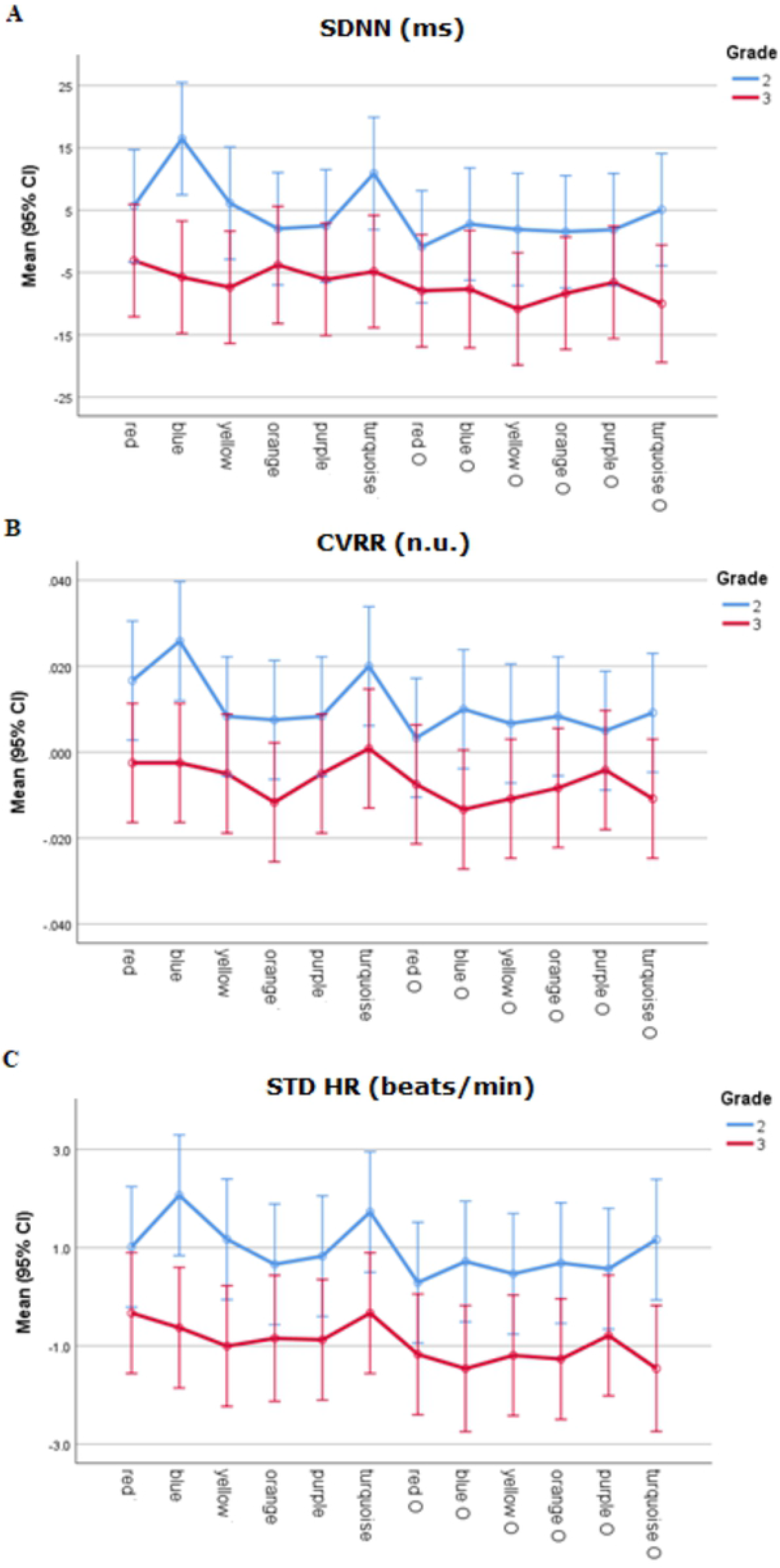
SDNN, CVRR and STD HR by grade and colour (normalized on white colour) Given that there were no systematic differences across colours in Table 3 for Reading Duration, Eye-tracking, EEG and EDA parameters, for a subsequent analysis all twelve background and overlay colours were averaged together in order to examine differences between younger and older children.

Grade comparisons (second vs. third) regarding the examined parameters over averaged scores for all colours together are presented in Table 4. A significant difference has been obtained regarding Reading duration, median Alpha, Betha, Delta, Theta power bands as well as for the Whole range, then for Fixation Duration Total and Fixation Duration Average and EDA. In all other parameters we observed no significant difference between second and third graders.

**Table 4.** Reading duration, EEG, Eye tracking and EDA parameters in second and third grade across all colours together - significant p values are marked in bold

| Parameters | Grade |  | p value* |
| --- | --- | --- | --- |
|  | Second (n=156) | Third (n=156) |  |
| Reading duration |  |  |  |
| RD (s) | 42.74 ± 31.81 | 50.33 ± 27.40 | 0.001 |
| EEG parameters (median power band) |  |  |  |
| Alpha | 13.50 ± 5.28 | 4.67 ± 3.20 | 0.001 |
| Beta | 6.36 ± 2.90 | 3.71 ± 4.70 | 0.001 |
| Delta | 71.49 ± 37.33 | 51.23± 48.90 | 0.001 |
| Theta | 17.55± 7.88 | 6.38± 4.75 | 0.001 |
| Whole Range | 123.36 ± 54.24 | 74.45± 61.30 | 0.001 |
| Eye tracking parameters |  |  |  |
| Fixation Count | 35.14 ± 13.97 | 36.35± 10.31 | 0.412 |
| Fixation Frequency [count/s] | 1.10 ± 0.89 | 1.01 ± 0.75 | 0.372 |
| Fixation Duration Total [s] | 40.01 ± 30.71 | 47.47±25.63 | 0.028 |
| Fixation Duration Average [ms] | 1038.49 ± 593.15 | 1283.29 ± 584.55 | 0.001 |
| Saccade Count | 31.31 ± 12.36 | 30.93 ± 8.05 | 0.753 |
| Saccade Frequency [count/s] | 0.99 ± 1.00 | 0.89 ± 0.71 | 0.376 |
| Saccade Duration Total [ms] | 662.9±293.4 | 712.0±273.6 | 0.147 |
| Saccade Duration Average [ms] | 21.65 ± 6.09 | 22.69 ± 3.88 | 0.093 |
| EDA value |  |  |  |
| EDA (uS) | 10.13± 3.25 | 6.32 ± 3.95 | 0.001 |

\*independent sample t test

Regarding Reading duration, third graders took significantly longer time to complete the reading. All EEG power bands (Alpha, Betha, Delta, Theta and Whole range) as well as EDA were higher in the second grade group, compared to third graders. An opposite trend was found for the eye-tracking measures whereby both Fixation Duration Total and Fixation Duration Average were higher in the third graders in comparison to the second graders.

## Discussion

We have evaluated differences in Reading duration, EEG, eye tracking, EDA and HRV parameters in 24 children (12 second and 12 third grade students of elementary school) using simultaneously monitored sensor signals.

The literature regarding EEG power bands (Alpha, Beta, Delta, Theta) has been shown to have a general developmental decreasing trend with an increasing age [23–25]. However, in the case of a specific mental activity, such as reading in this study, little is yet known about developmental changes of contribution of EEG power bands [25]. When kids were given a task to read on a white colour (which is a standard and everyday background colour), they showed a decrease in Alpha, Beta and Theta power bands with age, Table 2. Given that this is a pilot study in this sense, it remains to be resolved whether this is specifically a developmental trend, or one that has to do with experience, or both in combination. To fully resolve this issue, one would also need to have a much wider range of age groups tested.

Correlation between HRV parameters and age in infants and childhood, caused by progressive maturation of the autonomous nervous system, is known in the literature [26–29]. However, the age impact on HRV parameters is more extensive in infancy and the early childhood period (up to eight years) [26]. In this study, participants were 8 and 9 years old, so they were in years of life when the age impact was much less significant on HRV parameters [30]. Consequently, the level of stress in our study was expected (and found) to be lower in the group with a higher strain [31]. Namely, SDNN, CVRR and STD HR were found to be lower in younger children when reading on white background, Table 2. However, we would not predominantly attribute these findings to the developmental causal factor, but at least to some extent to experience in reading. This study gives further support to existing findings that colours may play an important role in the reading process [13,15,16,18,32–36]. When normalization on the white background was performed (by subtracting white from each of the other background and overlay colours), systematic differences (on at least two colours) were found regarding HRV parameters (normalized SDNN, CVRR, and STD HR values), with second graders scoring higher on these parameters, Table 3. The corresponding graphics across colours are presented in Figure 2, where it could be observed that blue and turquoise backgrounds have a calming impact (increasing normalized SDNN, CVRR, and STD HR values) on second graders, which is in accordance with the previous reports [37–39]. No systematic differences were found across colours between younger and older children in Reading duration, EEG power band, Eye-tracking and EDA measures. This is why, based on the results presented in Table 3, for the following analyses all twelve background and overlay colours were averaged together and examined differences between second and third graders in that case, Table 4. Based on the results presented in Table 4, we observed a significant difference across multimodal sensor measurements for reading on background and overlay colours in contrast to reading on the white background colour.

The findings concerning Reading duration on background and coloured overlays showed that the third-grade students have longer Reading duration in comparison to the second-graders. This could be explained by the fact that third graders or older children need more time to adapt to unexpected text and stimuli such as colour [40–42]. Also, previous studies have shown that older children are slower in reading than younger children because they make longer fixation duration and saccades [43–48] skipping words more frequently than younger [49,50]. Concerning the eye-tracking measures, it was in fact found that third-graders have longer Fixation Duration Total and Fixation Duration Average in comparison to the second-graders, which is a result that is in line with the above mentioned assertion that older children take longer to read. This indicates higher mental load in the reading task in older children.

A similar pattern of EEG results found for the white background was also found for the background and overlay colours. However, the number of parameters which showed a significant difference between second and third graders and significance levels were much more prominent for background and overlay colours in comparison to the white background. The study shows significant differences between the second and third graders in all four EEG power bands (Alpha, Beta, Delta and Theta), as well as in the Whole Range based on the averaged results from the twelve background and overlay colours.

Similarly, it is important to mention that EDA linearly correlates to arousal and reflects cognitive activity and emotional response [51,52] and it is the most used psychophysiological measure of arousal [53]. The higher the arousal is, the higher electrodermal activity is. In the present study, it was found to be higher among second graders, from which we conclude that they had a higher stress-level.

## Conclusion

The aim of this study was to assess developmental differences in second and third grade students of elementary school regarding reading on white vs coloured overlay and background and to evaluate differences in reading duration, brain, eye, electrodermal and heart activities. Evaluating all findings and results during reading on white colour and all 12 background and overlay colours, we can conclude that there is a decreasing trend with age regarding EEG power bands (Alpha, Beta, Delta, Theta) that is shown in the comparison between second and third grade students. In addition, second graders show lower scores of reading duration and eye-tracking measures (fixation duration total and fixation duration average), which confirms the fact that older children need more time to adapt on unexpected text and have longer fixations on words during reading. Comparing HRV parameters in second and third graders during reading on white colour, we have found lower scores in second graders and higher scores in 12 overlays and background colours compared to third graders, especially for SDNN, CVRR, and STD HR measures. The highest values of normalized SDNN, CVRR, and STD HR among students were reached in turquoise and blue background colours, which could be the result of calming effect during task performance caused by the background colour. Furthermore, we have found that EDA linearly correlates to level of arousal (tension and stress), where we have found higher values in younger children. Across single colours and its influence on measures during the reading task, there is no systematic difference in Reading duration, EEG power band, Eye-tracking and EDA measures, except for HRV measures as mentioned above.

Thus, the goal of combining different modalities was to find a more objective approach to understanding the developmental differences in children’s reading as well as to understand the contribution of different modalities and combinations of modalities in the process of reading text on a white vs colour overlay and background.

In the following work, it will be necessary to move forward from group studies to individual studies in order to determine and establish individual optimal parameters, as well as colours corresponding to individual differences in the reading process.

## Acknowledgment

The authors acknowledge the financial support from the Slovenian Research Agency (research core funding No. P2-0098), AD Futura Found (Public Scholarship, Development, Disability and Maintenance Found of the Republic of Slovenia), IPS Jozef Stefan and the Ministry of Education, Science and Technological Development of the Republic of Serbia.

## References

1. Schroeder S, Hyönä J, Liversedge SP. Developmental eye-tracking research in reading: Introduction to the special issue [Internet]. Vol. 27, Journal of Cognitive Psychology. Psychology Press Ltd; 2015 [cited 2020 Jul 11]. p. 500–10. Available from:https://www.tandfonline.com/doi/abs/10.1080/20445911.2015.1046877

2. Korneev AA, Matveeva EY, Akhutina T v. What We Can Learn about Reading Development from the Analysis of Eye Movements. Human Physiology. 2018 Mar 1;44(2):183–90.

3. Lobier M, Dubois M, Valdois S. The Role of Visual Processing Speed in Reading Speed Development. Barton JJS, editor. PLoS ONE [Internet]. 2013 Apr 4 [cited 2020 Mar 30];8(4):e58097. Available from: http://dx.plos.org/10.1371/journal.pone.0058097

4. Lerkkanen MK, Rasku-Puttonen H, Aunola K, Nurmi JE. Reading performance and its developmental trajectories during the first and the second grade. Learning and Instruction. 2004 Apr 1;14(2):111–30.

5. Miller B, Shriver EK, O’donnell C. Opening a Window into Reading Development: Eye Movements’ Role Within a Broader Literacy Research Framework. Vol. 42, School Psych Rev. 2013.

6. Korneev AA, Akhutina T v, Matveeva EYu. Reading in third graders with different state of the skill: an eye-tracking study. Moscow University Psychology Bulletin. 2019;2:64–87.

7. Vorstius C, Radach R, Lonigan CJ. Eye movements in developing readers: A comparison of silent and oral sentence reading. Visual Cognition. 2014;22(3):458–85.

8. Hulme C, Snowling MJ. Learning to Read: What We Know and What We Need to Understand Better.

9. McArthur G, Castles A. Helping children with reading difficulties: some things we have learned so far. npj Science of Learning [Internet]. 2017 Dec 31 [cited 2020 Jul 11];2(1):7. Available from:www.motif.org.au

10. van Bommel WJM, van den Beld GJ. Lighting for work: A review of visual and biological effects. Lighting Research and Technology [Internet]. 2004 Dec 19 [cited 2020 Mar 4];36(4):255–69. Available from: http://journals.sagepub.com/doi/10.1191/1365782804li122oa

11. de Jong PF, van der Leij A. Effects of Phonological Abilities and Linguistic Comprehension on the Development of Reading. Scientific Studies of Reading. 2002 Jan;6(1):51–77.

12. Uccula A, Enna M, Mulatti C. Colors, colored overlays, and reading skills. Frontiers in Psychology [Internet]. 2014 Jul 29 [cited 2020 Jul 10];5(JUL):833. Available from:http://journal.frontiersin.org/article/10.3389/fpsyg.2014.00833/abstract

13. Conway ML, Evans BJW, Evans JC, Suttle CM, Engel FL, Child PBASIC, et al. Colors, colored overlays, and reading skills. Allen P, editor. Frontiers in Psychology [Internet]. 2014 Sep 20 [cited 2019 Jun 10];5(7):9–21. Available from: www.frontiersin.org

14. Jansky JJ. A case of severe dyslexia with aphasic-like symptoms. Bulletin of the Orton Society [Internet]. 1958 May [cited 2020 May 5];8(1):8–11. Available from:http://link.springer.com/10.1007/BF02657600

15. Wilkins AJ, Evans BJW. Visual stress, its treatment with spectral filters, and its relationship to visually induced motion sickness. Applied Ergonomics [Internet]. 2010 Jul [cited 2020 Mar 3];41(4):509–15. Available from:https://linkinghub.elsevier.com/retrieve/pii/S0003687009000325

16. Pinna B, Deiana K. On the Role of Color in Reading and Comprehension Tasks in Dyslexic Children and Adults. i-Perception [Internet]. 2018 May 9 [cited 2020 Apr 10];9(3):204166951877909. Available from:http://journals.sagepub.com/doi/full/10.1177/2041669518779098

17. Denton TF, Meindl JN. The Effect of Colored Overlays on Reading Fluency in Individuals with Dyslexia. Behavior Analysis in Practice [Internet]. 2016 Sep [cited 2020 Jul 10];9(3):191–8. Available from: /pmc/articles/PMC4999357/?report=abstract

18. Veszeli J, Shepherd AJ. A comparison of the effects of the colour and size of coloured overlays on young children’s reading. Vision Research. 2019 Mar 1;156:73–83.

19. Sam W. The use of eye tracking with infants and children. Practical Research With Children. Routledge. 2016; pp.24–45.

20. Giagloglou E, Radenkovic M, Brankovic S, Antoniou P, Zivanovic-Macuzic I. Pushing, pulling and manoeuvring an industrial cart: a psychophysiological study. International Journal of Occupational Safety and Ergonomics [Internet]. 2019 Apr 3 [cited 2020 Jul 13];25(2):296–304. Available from: https://pubmed.ncbi.nlm.nih.gov/28849989/

21. Sachin S, Netaji G N. Pattern analysis of different ECG signal using Pan-Tompkin’s algorithm. (IJCSE) International Journal on Computer Science and Engineering. 2010.

22. Shaffer F, Ginsberg JP. An Overview of Heart Rate Variability Metrics and Norms. Frontiers in Public Health [Internet]. 2017 Sep 28 [cited 2020 Jul 11];5:258. Available from: /pmc/articles/PMC5624990/?report=abstract

23. Marshall PJ, Bar-Haim Y, Fox NA. Development of the EEG from 5 months to 4 years of age. Clinical Neurophysiology. 2002 Aug 1;113(8):1199–208.

24. Miskovic V, Ma X, Chou C-A, Fan M, Owens M, Sayama H, et al. Developmental changes in spontaneous electrocortical activity and network organization from early to late childhood. NeuroImage [Internet]. 2015 Sep 1 [cited 2020 May 24];118:237–47. Available from:https://linkinghub.elsevier.com/retrieve/pii/S1053811915005108

25. Spironelli C, Angrilli A. Developmental aspects of language lateralization in delta, theta, alpha and beta EEG bands. Biological Psychology [Internet]. 2010 Oct [cited 2020 Jul 12];85(2):258–67. Available from: https://pubmed.ncbi.nlm.nih.gov/20659528/

26. Massin M, von Bernuth G. Normal ranges of heart rate variability during infancy and childhood. Pediatric Cardiology. 1997 Jul;18(4):297–302.

27. Goto M, Nagashima M, Baba R, Nagano Y, Yokota M, Nishibata K, et al. Analysis of heart rate variability demonstrates effects of development on vagal modulation of heart rate in healthy children. Journal of Pediatrics. 1997;130(5):725–9.

28. Finley JP, Nugent ST. Heart rate variability in infants, children and young adults. Journal of the Autonomic Nervous System [Internet]. 1995 Feb 9 [cited 2020 May 25];51(2):103–8. Available from: https://linkinghub.elsevier.com/retrieve/pii/0165183894001173

29. Seppälä S, Laitinen T, Tarvainen MP, Tompuri T, Veijalainen A, Savonen K, et al. Normal values for heart rate variability parameters in children 6-8 years of age: the PANIC Study. Clinical Physiology and Functional Imaging [Internet]. 2014 Jul [cited 2020 May 25];34(4):290–6. Available from: http://doi.wiley.com/10.1111/cpf.12096

30. Kim HG, Cheon EJ, Bai DS, Lee YH, Koo BH. Stress and heart rate variability: A meta-analysis and review of the literature [Internet]. Vol. 15, Psychiatry Investigation. Korean Neuropsychiatric Association; 2018 [cited 2020 Jul 12]. p. 235–45. Available from: /pmc/articles/PMC5900369/?report=abstract

31. Kang MG, Koh SB, Cha BS, Park JK, Woo JM, Chang SJ. Association between job stress on heart rate variability and metabolic syndrome in shipyard male workers. Yonsei Medical Journal. 2004 Oct 31;45(5):838–46.

32. Freiders S, Lee S, Statz D, Kim Group T. The Influence of Color on Physiological Response.

33. Morrison R. Effect of Color Overlays on Reading Efficiency [Internet]. 2011 [cited 2020 Apr 10]. Available from: https://scholarworks.umass.edu/open_access_dissertations/431

34. Griffiths PG, Taylor RH, Henderson LM, Barrett BT. The effect of coloured overlays and lenses on reading: a systematic review of the literature. Ophthalmic and Physiological Optics [Internet]. 2016 Sep 1 [cited 2020 Apr 10];36(5):519–44. Available from:http://doi.wiley.com/10.1111/opo.12316

35. Rello L, Bigham JP. Good Background Colors for Readers: A Study of People with and without Dyslexia. 2017 [cited 2020 Apr 10]; Available from: https://doi.org/10.1145/3132525.3132546

36. Hlengwa N, Moonsamy P, Ngwane F, Nirghin U, Singh S. The effect of color overlays on the reading ability of dyslexic children [Internet]. Vol. 65, Indian Journal of Ophthalmology. Medknow Publications; 2017 [cited 2020 Jul 12]. p. 772–3. Available from:https://www.ncbi.nlm.nih.gov/pmc/articles/PMC5598199/

37. AL-Ayash A, Kane RT, Smith D, Green-Armytage P. The influence of color on student emotion, heart rate, and performance in learning environments. Color Research & Application [Internet]. 2016 Apr 1 [cited 2020 May 24];41(2):196–205. Available from:http://doi.wiley.com/10.1002/col.21949

38. Mehta R, Zhu R. Blue or red? Exploring the effect of color on cognitive task performances. Science [Internet]. 2009 Feb 27 [cited 2020 Mar 4];323(5918):1226–9. Available from:https://www.sciencemag.org/lookup/doi/10.1126/science.1169144

39. Moharreri S., Dabanloo N. J., Parvaneh S., & Nasrabadi A. M. How to interpret psychology from heart rate variability? 2011 1st Middle East Conference on Biomedical Engineering. 2011. doi:10.1109/mecbme.2011.5752124

40. Rayner K, Liversedge SP, White SJ. Eye movements when reading disappearing text: The importance of the word to the right of fixation. Vision Research. 2006 Feb 1;46(3):310–23.

41. Rayner K, Yang J, Schuett S, Slattery TJ. Eye movements of older and younger readers when reading unspaced text. Experimental psychology. 2013;60(5):354–61.

42. Rayner K, Castelhano MS, Yang J. Eye Movements and the Perceptual Span in Older and Younger Readers. Psychology and Aging. 2009.

43. Kliegl R, Grabner E, Rolfs M, Engbert R. Length, frequency, and predictability effects of words on eye movements in reading. In: European Journal of Cognitive Psychology. 2004. p. 262–84.

44. Dambacher M, Kliegl R, Hofmann M, Jacobs AM. Frequency and predictability effects on event-related potentials during reading. Brain Research. 2006;1084(1):89–103.

45. Rayner K, Castelhano MS, Yang J. Preview benefit during eye fixations in reading for older and younger readers. Psychology and Aging. 2010;25(3):714–8.

46. Rayner K, Chace KH, Slattery TJ, Ashby J. Eye movements as reflections of comprehension processes in reading. Vol. 10, Scientific Studies of Reading. 2006;p. 241–55.

47. Rayner K. Eye movements and the perceptual span in beginning and skilled readers. Journal of Experimental Child Psychology. 1986;41(2):211–36.

48. Rayner K, Foorman BR, Perfetti CA, Pesetsky D, Seidenberg MS. How Psychological Science Informs the Teaching of Reading. Psychological Science in the Public Interest. 2001;2(2):31–74.

49. Laubrock J, Kliegl R, Engbert R. SWIFT explorations of age differences in eye movements during reading. Vol. 30, Neuroscience and Biobehavioral Reviews. 2006; p. 872–84.

50. Rayner K, Reichle ED, Stroud MJ, Williams CC, Pollatsek A. The effect of word frequency, word predictability, and font difficulty on the eye movements of young and older readers. Psychology and Aging. 2006;21(3):448–65.

51. Boucsein Wolfram. Electrodermal activity. Springer Science+Business Media, LLC. 2012; p. 618.

52. Boucsein W, Boucsein W. Principles of Electrodermal Phenomena. In: Electrodermal Activity. Springer US; 2012; p. 1–86.

53. Carlucci L, Watkins MW, Sergi MR, Cataldi F, Saggino A, Balsamo M. Dimensions of anxiety, age, and gender: Assessing dimensionality and measurement invariance of the State-Trait for Cognitive and Somatic Anxiety (STICSA) in an Italian sample. Frontiers in Psychology. 2018;p. 2345.

